# Drug Repurposing for the SARS-CoV-2 Papain-Like Protease

**DOI:** 10.1101/2021.06.04.447160

**Authors:** Chia-Chuan D. Cho, Shuhua G. Li, Kai S. Yang, Tyler J. Lalonde, Ge Yu, Yuchen Qiao, Shiqing Xu, Wenshe Ray Liu

**Affiliations:** The Texas A&M Drug Discovery Laboratory, Department of Chemistry, Texas A&M University, College Station, TX 77843, USA; Institute of Biosciences and Technology and Department of Translational Medical Sciences, College of Medicine, Texas A&M University, Houston, TX 77030, USA; Department of Biochemistry and Biophysics, Texas A&M University, College Station, TX 77843, USA; Department of Molecular and Cellular Medicine, College of Medicine, Texas A&M University, College Station, TX 77843, USA

**Keywords:** COVID-19, SARS-CoV-2, papain-like protease, deubiquitinase, cysteine protease

## Abstract

As the pathogen of COVID-19, SARS-CoV-2 encodes two essential cysteine proteases that process the pathogen’s two large polypeptide translates ORF1a and ORF1ab in human host cells to form 15 functionally important, mature nonstructural proteins. One of the two enzymes, papain-like protease or PLpro, also possesses deubiquitination and deISGylation activities that suppresses host innate immune responses toward SARS-CoV-2 infection. Therefore, PLpro is a potential COVID-19 drug target. To repurpose drugs for PLpro, we experimentally screened 33 deubiquitinase and 37 cysteine protease inhibitors on their inhibition of PLpro. Our results showed that 15 deubiquitinase and 1 cysteine protease inhibitors exhibit potent inhibition of PLpro at 200 *μ*M. More comprehensive characterizations revealed 7 inhibitors GRL0617, SJB2-043, TCID, DUB-IN-1, DUB-IN-3, PR-619, and S130 with an IC_50_ value below 60 *μ*M and four inhibitors GRL0617, SJB2-043, TCID, and PR-619 with an IC_50_ value below 10 *μ*M. Among four inhibitors with an IC_50_ value below 10 *μ*M, SJB2-043 is the most unique in that it doesn’t fully inhibit PLpro but has an outstanding IC_50_ value of 0.56 *μ*M. SJB2-043 likely binds to an allosteric site of PLpro to convene its inhibition effect, which needs to be further investigated. As a pilot study, the current work indicates that COVID-19 drug repurposing by targeting PLpro holds promises but in-depth analysis of repurposed drugs is necessary to avoid omitting allosteric inhibitors.

## INTRODUCTION

The current prevailing pandemic of COVID-19 (coronavirus disease 2019) has caused the global health and economic consequences that is often compared to that of 1918 influenza pandemic.^1, 2^ As of June 1^st^, 2021, the total number of confirmed global COVID-19 cases has exceeded 170 million, of which more than 3.5 million have succumbed to death (WHO data). Encouragingly, three COVID-19 vaccines developed by Pfizer/BioNtech, Moderna, and Johnson & Johnson have been approved by FDA for human immunization in the United States. Although vaccines are promising in containing the pandemic, their availability does not diminish the urgent need for other effective antiviral drugs. Existing COVID-19 vaccines are targeting the membrane Spike protein of SARS-CoV-2, the pathogen of COVID-19. Spike is highly mutable.^3^ New viral strains with critical mutations in Spike such as UK and South Africa SARS-CoV-2 strains have emerged. Effectiveness of vaccines against these new strains will be a concern.^3^ Vaccines are also preventative making them not an option for the treatment of COVID-19 patients. For quick access to effective antivirals for COVID-19, repurposing of existing drugs has been broadly conducted. This effort has led to the identifications of a number of potential antivirals for SARS-CoV-2.^4^ So far, remdesivir is the only antiviral that has been approved for the treatment of COVID-19. Remdesivir is a nucleoside analog that inhibits RNA-dependent RNA polymerase (RdRp) of SARS-CoV-2. It shows high potency in inhibiting SARS-CoV-2 *in vitro* but appears to be only modestly effective in treating COVID-19 patients.^5^ In view of the ongoing pandemic in its colossal scale and apocalyptic impact, there is a dire need of more effective antivirals with novel mechanisms of action to save lives of COVID-19 patients.

As the pathogen of COVID-19, SARS-CoV-2 has a positive-sensed genomic RNA.^6–8^ It encodes 10 open reading frames (ORFs). As the largest ORF, ORF1ab comprises more than two thirds of the whole genome. In an infected host cell, ORF1ab is translated to two large polypeptide translates, ORF1a (~500 kDa) and ORF1ab (~800 kDa),^9^ by the host protein translation system. ORF1ab is formed due to a frameshift during protein translation. Both ORF1a and ORF1ab polypeptides need to undergo proteolytic cleavage to form 15 mature proteins. These mature proteins are nonstructural proteins (Nsps) essential for the virus in its reproduction and pathogenesis. The proteolytic cleavage of ORF1a and ORF1ab is an autocatalytic process. Two internal polypeptide regions, nsp3 and nsp5, possess cysteine protease activity that cleaves themselves and all other nsps in ORF1a and ORF1ab. Nsp3 is commonly referred to as papain-like protease (PLpro) and nsp5 as main protease (Mpro).^10^ Both PLpro and Mpro are essential to SARS-CoV-2. Of the two enzymes, PLpro recognizes the tetrapeptide LXGG motif that is in-between viral proteins nsp1 and nsp2, nsp2 and nsp3, and nsp3 and nsp4. Its cleavage of peptide bonds leads to the release of nsp1, nsp2, and nsp3 which are essential for host modulation and viral replication.^11^ In addition, recent studies have shown that PLpro can proteolytically remove K48-crosslinked ubiquitin (Ub) and interferon-stimulated gene 15 product (ISG15), both of which play important roles in the regulation of host innate immune responses to viral infection.^12, 13^ PLpro is also conserved across various coronoviruses.^14^ For these reasons, PLpro is an attractive COVID-19 drug target for the development of antivirals that may be potentially used as broad-spectrum inhibitors for other coronaviruses as well.^15–19^

In view of the urgent need of antivirals for effective COVID-19 treatments, repurposing approved drugs and late-stage clinical drug candidates against SARS-CoV-2 is an efficient strategy. Significant progress has been made in drug repurposing for Mpro. The first orally administered SARS-CoV-2 Mpro inhibitor PF-07321332 that was developed by Pfizer is currently undergoing evaluation in a Phase 1b multi-dose study in hospitalized COVID-19 participants. In contrast, inhibitors for PLpro remain relatively underdeveloped because PLpro is comparably a more challenging drug target. It has a flatter active site pocket than Mpro. Since PLpro is both a cysteine protease and a deubiquitinase, we reasoned that screening of cysteine protease inhibitors and deubiquitinase inhibitors would provide fast identification of PLpro inhibitors as initial hits for further optimization. In this study, we report our progress in identifying PLpro inhibitors by screening 33 deubiquitinase inhibitors and 37 cysteine protease inhibitors. Among these molecules, we identified seven drug candidates that can potently inhibit PLpro with an IC_50_ value below 60 μM. Four molecules have an IC_50_ value below 10 μM. One molecule SBJ2-043 can only partially inhibit PLpro but has a remarkable low IC_50_ value of 0.56 μM. More in-depth investigation of these inhibitors in their mechanisms of action will likely reveal unique inhibition mechanisms for PLpro for more advanced development of potent PLpro inhibitors.

## RESULTS & DISCUSSION

PLpro is a cysteine protease with a classical Cys-His-Asp catalytic triad (Cys111, His272 and Asp286). It is responsible for processing three cleavage sites in the viral polypeptides ORF1a and ORF1ab to release nsp1, nsp2, and nsp3. Apart from proteolytic processing, PLpro has also deubiquitinase and deISGylase activities to help SARS-CoV-2 evade host immune response.^12, 13, 20–22^ Thus, to identify new PLpro inhibitors, we screened 33 deubiquitinase inhibitors and 37 cysteine protease inhibitors on their inhibition of PLpro.

To express PLpro for experimental characterizations of selected small molecules, we constructed an PLpro vector for expression in *E. coli* (Figure 1A). In this construct, PLpro was fused to the *C* terminus of superfolder green fluorescent protein (sfGFP) that is known to stabilize a fused partner.^23^ A 6×His tag was fused to N terminus of sfGFP for affinity purification using Ni-NTA resins. Between sfGFP and PLpro was a TEV cleavage site for the proteolytic removal of PLpro from the fusion protein by TEV protease. During the purification and treatment of the sfGFP-PLpro fusion, we noticed that the cleaved PLpro quickly aggregates. Therefore, for long term storage of PLpro, we have chosen to save and use sfGFP-PLpro instead of PLpro for all our assays. In order to test catalytic activities of PLpro, we synthesized a fluorogenic protein substrate Ub-AMC (AMC: 7-amino-4-methylcoumarin) using our recently developed activated cysteine-directed protein ligation technique and purchased a fluorogenic peptide substrate Z-LRGG-AMC (Figure 1B).^24^ The hydrolysis of Ub-AMC and Z-LRGG-AMC releases AMC whose strong blue fluorescence can be traced using a fluorometer or a fluorescent plate reader. To obtain an optimal assay condition for inhibitor characterization, both Ub-AMC and Z-LRGG-AMC are titrated at 20 nM PLpro (Figure 1C and 1D). Based on the product formation curves, we deemed that 5 μM Ub-AMC and 50 μM Z-LRGG-AMC provide easily detectable linear production formation within 1000 s and therefore used these two conditions in our following inhibitor characterization assays. We have also characterized the influence of DMSO to the PLpro catalysis since most commercial small molecules are provided as 10 mM DMSO stocks. The stability of PLpro in the presence of DMSO restricts the highest drug concentration we can test. We titrated DMSO from 0.1% to 20% (Figure 1E). It showed that PLpro had slightly reduced activity at 2% DMSO and this inhibition trend by DMSO increased when the DMSO concentration improves. Considering most of commercial small molecules provided as 10 mM DMSO stocks, it will be 200 μM in an assay when it is finally diluted by 50 folds as in 2% DMSO. At this condition, the PLpro remains almost 100% activity and is good for quantifying drug inhibition effects.

**Figure 1.**
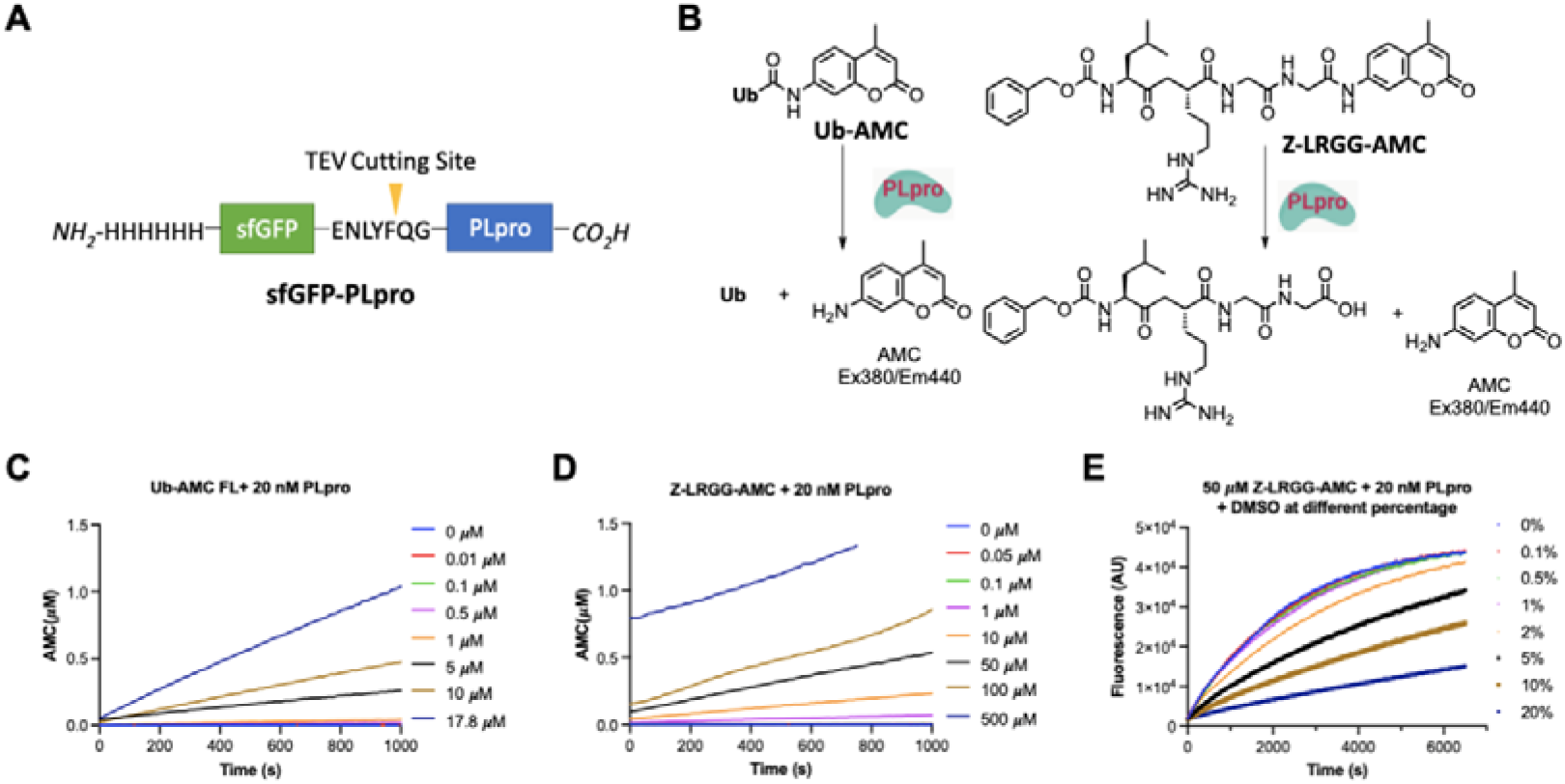
(A) The structures of the sfGFP-PLpro fusion protein whose expression in *E. coli* have been tested. (B) Structures of two PLpro substrates and their catalytic release of AMC. (C) The catalytic release of AMC at various concentrations of Ub-AMC by 20 nM PLpro. (D) The catalytic release of AMC at various concentrations of Z-LRGG-AMC by 20 nM PLpro. (E) The catalytic release of AMC at 50 μM LRGG-AMC by 20 nM PLpro in the presence of various concentrations of DMSO.

Since PLpro is a cysteine protease that is also a deubiquitinase, we initiated our search of its inhibitors using 31 deubiquitinase inhibitors (Figure 2) and 37 cysteine protease inhibitors (Figure 3) that we purchased from MedChemExpress. These molecules were provided as 10 mM DMSO stocks without further purification. Two additional deubiquitinase inhibitors C527 and VLX1570 were acquired from Cayman Scientific (Figure 2). As an initial screening assay, all acquired small molecules were analyzed at a 200 μM final concentration. In this assay, an inhibitor (400 μM) was incubated with 40 nM PLpro, 4 mM DTT, and 4% DMSO in a reaction buffer containing 20 mM Tris and 300 mM NaCl at pH 7.5 for 30 min. 100 μL of this incubation solution was then mixed with an equal volume of a substrate solution containing 100 μM Z-LRGG-AMC in the reaction buffer to initiate the catalytic release of AMC. The AMC fluorescence (Ex380/Em440) was recorded in a plate reader and the linear slope within the first 10 min of a reaction was calculated as the initial product formation rate. Among the deubiquitinase inhibitors at 200 μM, GRL0617 and TCID inhibited PLpro activity almost completely, PR-619 and ML364 had significant quenching effects leading to negative signals detected for the two compounds, and SjB2-043, P22077, LDN-57444, HBX19818, NSC632839, P005091, GSK2643943A and SJB3-019A inhibited PLpro partially but at significant levels (Figure 4A). In contrast, among 37 cysteine protease inhibitors, only S130 displayed strong inhibition of PLpro at the 200 μM level (Figure 4B).

**Figure 2.**
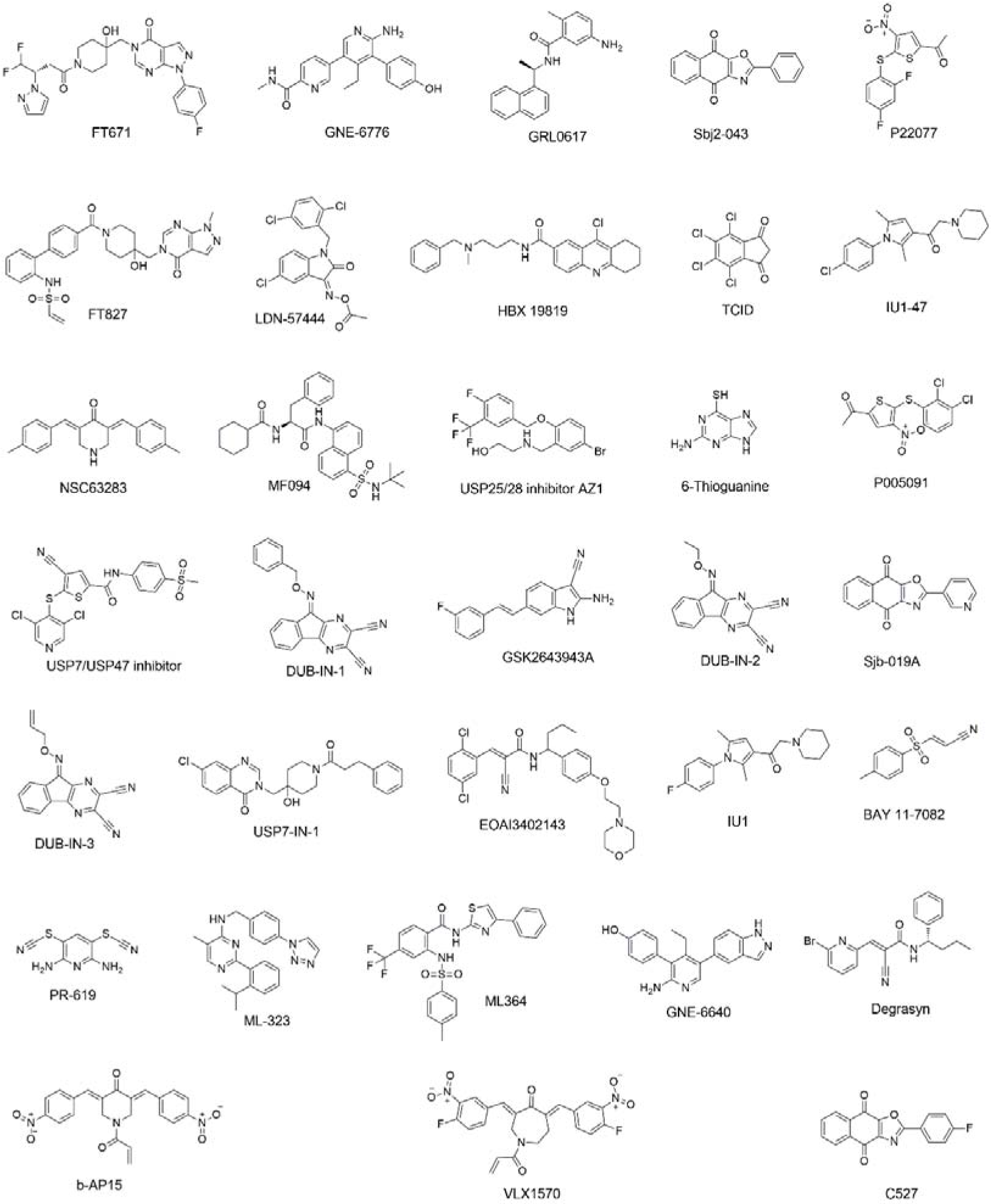
Structures of 33 selected deubiquitinase inhibitors.

**Figure 3.**
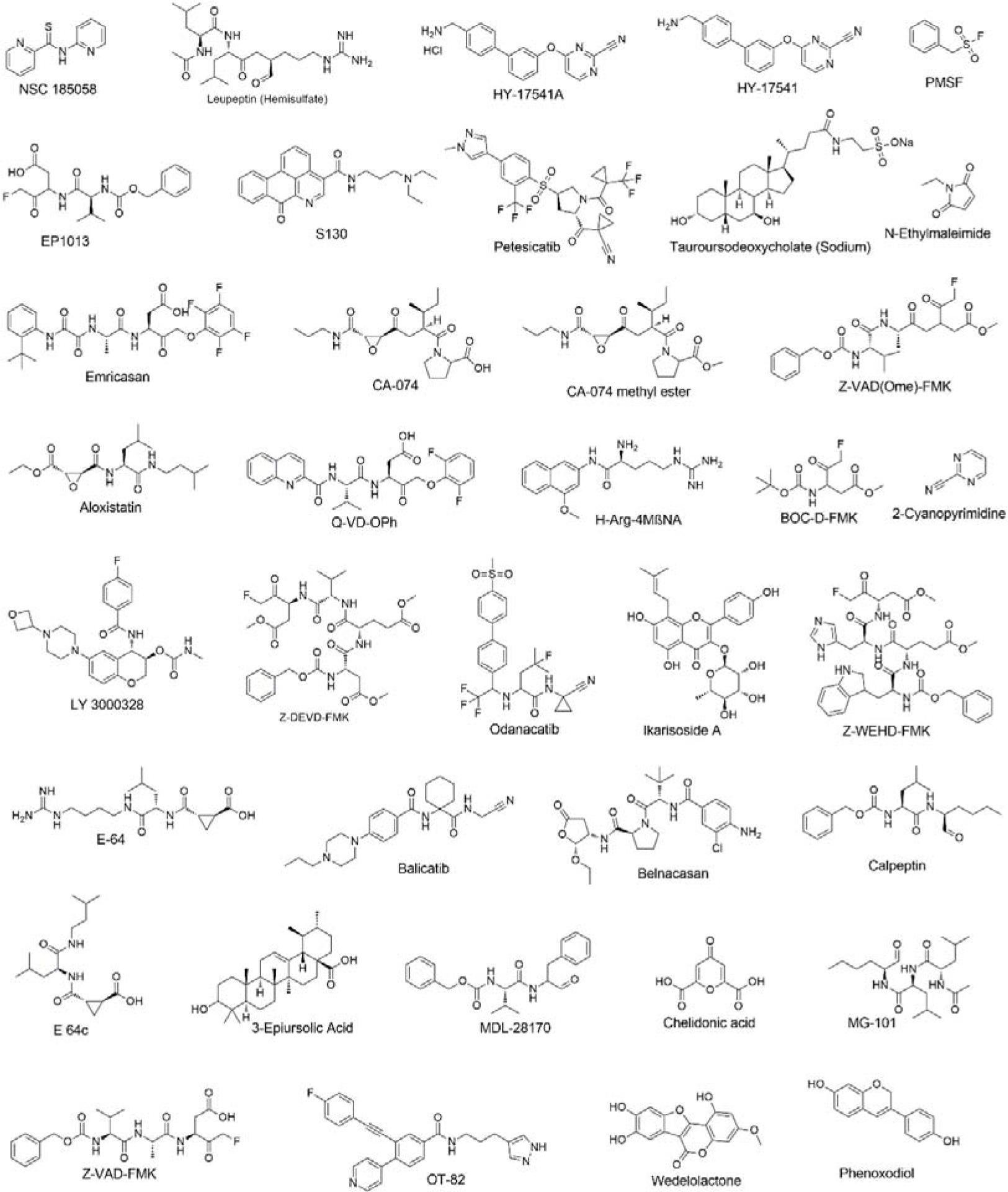
Structures of 37 selected cysteine protease inhibitors.

**Figure 4.**
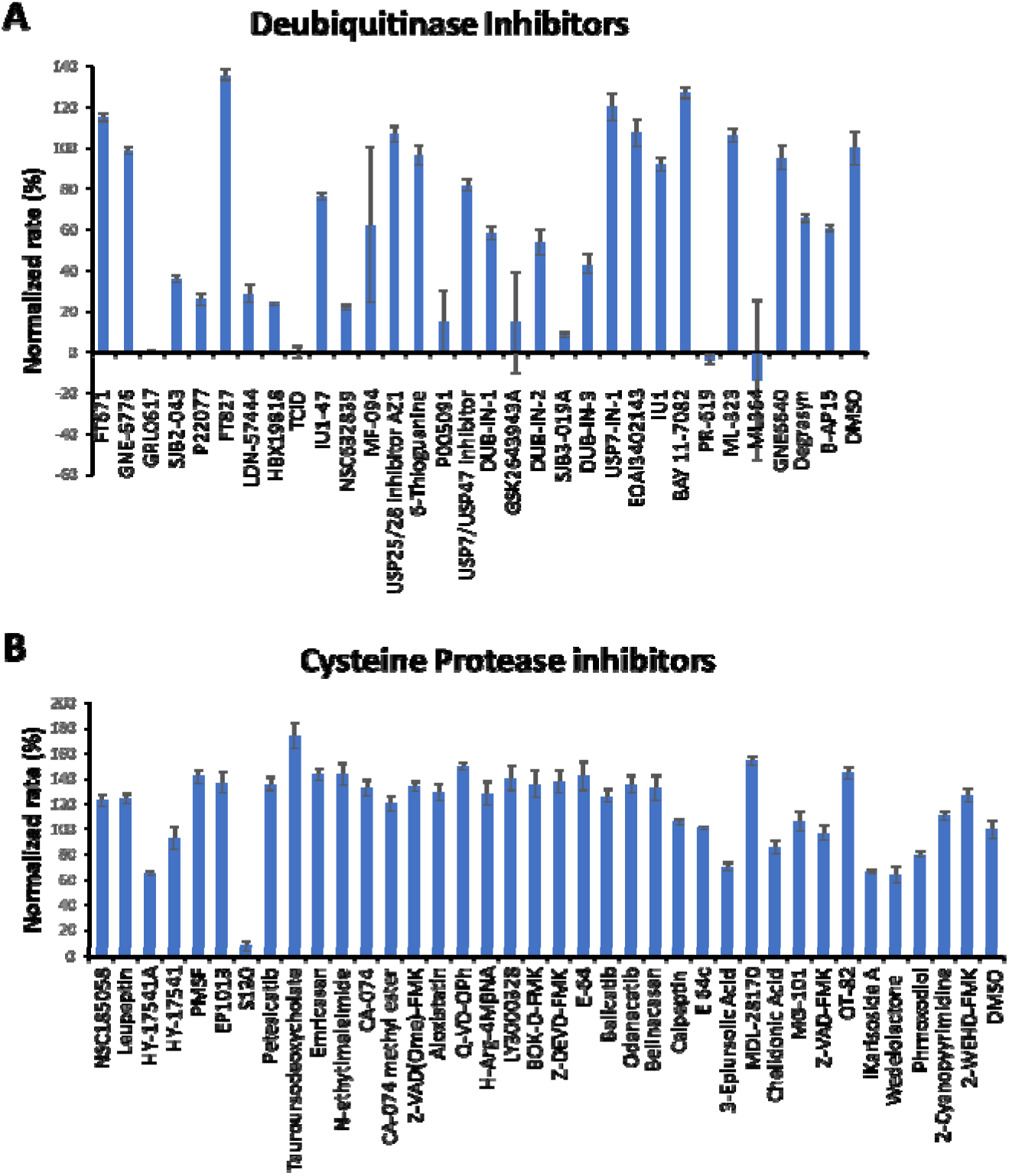
(A) Initial screening of PLpro inhibition by 33 deubiquitinase inhibitors. (B) Initial screening of PLpro inhibition by 37 cysteine protease inhibitors. Reaction conditions: 20 nM PLpro, 50 μM Z-LRGG-AMC, 200 μM inhibitor, 20 mM Tris-HCl, 300 mM NaCl, pH 7.5.

For 16 small molecules that exhibited significant inhibition of PLpro at the 200 μM level (PR-619 and ML364 were selected as well), we further conducted characterizations of their IC_50_ values. For IC_50_ characterizations, AMC product formation rates at 20 nM PLpro, 50 μM Z-LRGG-AMC and varied concentrations of a small molecule were measured in triplicates. Data of AMC product formation rate vs small molecule concentration were fitted nonlinearly to a sigmodal inhibition equation to determine IC_50_ values in GraphPad Prism 9. All data are presented in Figure 5. Except for PR-619 and ML364, we adjusted the initial rate by subtracting autofluorescence and adding photobleaching from a provided inhibitor. After this adjustment, we noticed that ML364 did not show strong concentration-dependent inhibition of PLpro but PR-619 strongly inhibited PLpro at concentration above 1 μM. Among all tested compounds, seven had a determined IC_50_ value below 60 μM. These included SJB2-043 (0.56 μM), GRL0617 (1.4 μM), PR-619 (6.1 μM), TCID (6.4 μM), SJB3-019A (8.15 μM), DUB-IN-3 (12.5 μM), and S130 (35.0 μM). GRL0617, TCID, PR-619 and S130 can fully inhibit PLpro at 200 μM. SJB2-043, SJB3-019a and DUB-IN-3 achieved only partial inhibition of PLpro at the highest tested concentration of 200 μM. GRL0617 is a naphthalene-based compound that was previously shown as a non-covalent inhibitor of PLpro from SARS-CoV with no significant inhibition of host proteases.^25, 26^ PLpro proteins from SARS-CoV and SARS-CoV-2 share 83% sequence identity and have a very high level of sequence and structural similarity in their substrate binding pockets.^13, 19, 27, 28^ In line with other recent studies, our results show that GRL0617 is also effective at inhibiting PLpro from SARS-CoV-2 (IC_50_=1.4 μM).^12, 27, 29^ Given its strong potency and relatively small size, structure-activity relationship studies of GRL0617 will likely lead to more potent PLpro inhibitors with high antiviral activities. TCID is an inhibitor for ubiquitin C-terminal hydrolase L3 with a reported IC_50_ of 0.6 μM.^30^ TCID has two ketones conjugated to a highly electron withdrawing tetrachlorobenzene. Both ketones are highly prone to hydrate or react with a nucleophilic residue in PLpro. It will be interesting to study how this molecule interacts with PLpro covalently or noncovalently. A structural investigation of its complex with PLpro will serve as a starting point for the development of more potent inhibitors for PLpro. PR619 is a thiocyanate reagent that broadly inhibits a series of deubiquitinases.^31^ The propensity of exchanging the cyano group with a protein thiolate makes PR619 likely to target cysteine proteases broadly. How exactly PR619 inactivates PLpro needs to be investigated further though a possible mechanism is the covalent transfer of the cyano group to the active site cysteine. S130 is an inhibitor that was originally discovered as an ATG4B inhibitor.^32^ ATG4B functions similar to a deubiquitinase and hydrolyzes the preprotein of ATG8. Since mutating the catalytic cysteine in ATG4B does not significantly affect the binding of S130 to ATG4B, S130 likely interacts with ATG4B noncovalently. A similar mechanism to inhibit PLpro is also expected. S130 has a large aromatic moiety in which four rings are conjugated. It will be interesting to see how this extended large aromatic moiety interacts with PLpro to exert strong binding. SJB2-043, SJB3-019a and DUB-IN-3 are three molecules that inhibit PLpro only partially at the highest tested concentration of 200 μM. SJB2-043 and SJB3-019a are two structurally similar compounds that inhibit USP1.^33^ Both have two quinone oxygens that potentially interact with nucleophilic residues in PLpro. Since both molecules did not inhibit PLpro completely but displayed clearly measurable IC_50_ values, they likely bind to an allosteric site of PLpro. Further investigations of these two molecules in their mechanisms of inhibiting PLpro will likely reveal novel targeting sites for the development of PLpro inhibitors. SJB2-043 has the lowest IC_50_ value among all tested small molecules, indicating very strong binding to PLpro. A potential application of this strong binding is to use it to develop a proteasome targeting chimera for PLpro. This is a direction we are actively exploring. DUB-IN-3 is a USP8 inhibitor that has two nitrile groups conjugated to a diazine.^34^ The electron deficient nature of the diazine makes the two nitriles reactive toward nucleophilic residues in PLpro. It is possible that one of these two nitriles will hit on the catalytic cysteine in PLpro to form a reversible covalent complex. The low Hill coefficient of the DUB-IN-3 inhibition curve also indicates a complicated inhibition mechanism.

**Figure 5.**
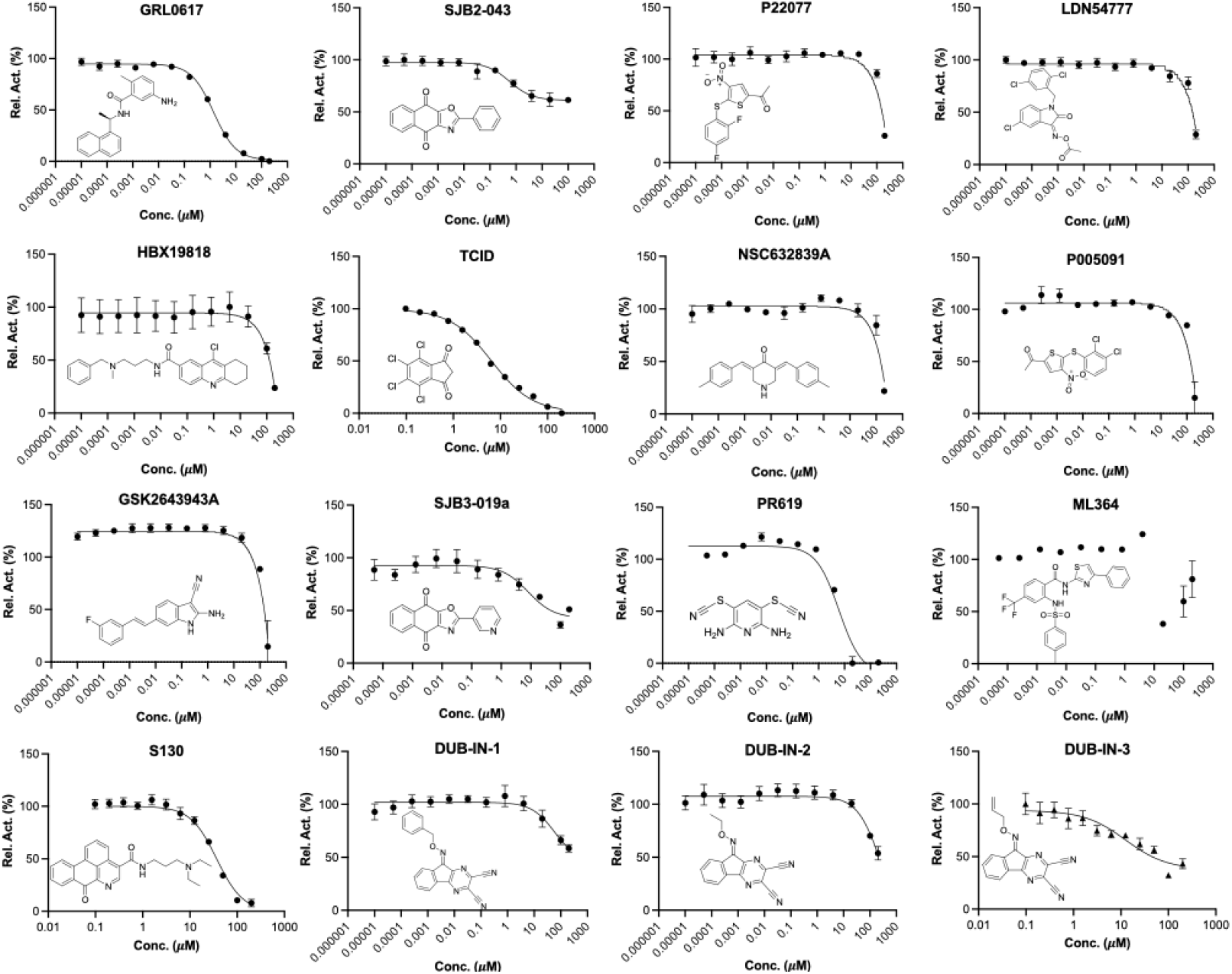
IC_50_ characterizations for 16 small molecules on their inhibition of PLpro using 50 μM LRGG-AMC as a substrate. Experiments at different conditions were performed in triplicates.

For six deubiquitinase inhibitors that showed high potency in inhibiting PLpro catalyzed AMC release from Z-LRGG-AMC, we have also characterized their inhibition of PLpro in its hydrolysis of Ub-AMC. Ub-AMC is much larger than Z-LRGG-AMC and involves a much bigger interface than Z-LRGG-AMC to interact with PLpro. For inhibitors that do not directly target the active site, they may display different inhibition characteristics when Ub-AMC instead of Z-LRGG-AMC is used as a substrate. We tested all six inhibitors in the presence of 20 nM PLpro and 5 μM Ub-AMC (Figure 6). When 5 μM Ub-AMC was used as a substrate, IC_50_ values of GRL0617, TCID and PR619 were determined as 1.80, 10.5 and 12.9 μM, respectively. These IC_50_ values are higher than but comparable to IC_50_ values determined when 50 μM Z-LRGG-AMC was used as a substrate, indicating that all three inhibitors are likely involved in a similar mechanism to inhibit PLpro. GRL0617 is known to bind the PLpro active site. Therefore, it is highly likely that TCID and PR619 bind to the PLpro active site as well. When Ub-AMC was used as a substrate, SJB2-043 had a determined IC_50_ value of 0.091 μM. This IC_50_ value is much lower than the determined IC_50_ value when Z-LRGG-AMC was used as a substrate. Similar to the observed pattern when Z-LRGG-AMC was used as a substrate, SJB2-043 did not inhibit PLpro completely. It is obvious that SJB2-043 behaves very differently from GRL0617, TCID and PR619. It binds likely to an allosteric site of PLpro. This allosteric binding influences apparently the catalytic hydrolysis of Ub-AMC more than Z-LRGG-AMC. This is likely attributed to the much larger binding interface of Ub-AMC that responses more sensitively to the PLpro structural perturbation. When Ub-AMC was used as a substrate, SBJ3-019a displayed an inhibition curve that is more complicated than that from SBJ2-043. It showed two inhibition stages with the first leading to partial inhibition and the second continuously inhibiting PLpro without reaching its inhibition plateau at the highest concentration we tested. Since it could not be fitted to a simple inhibition mechanism, we did not calculate its IC_50_ value. When Ub-AMC was used as a substrate, DUB-IN-3 displayed a simple inhibition curve that did not reach its plateau at the highest inhibitor concentration. Its estimated IC_50_ value was above 10 μM and similar to the IC_50_ value determined when Z-LRGG-AMC was used as a substrate. This similarity indicates that DUB-IN-3 likely binds to the PLpro active site to convene its inhibition.

**Figure 6.**
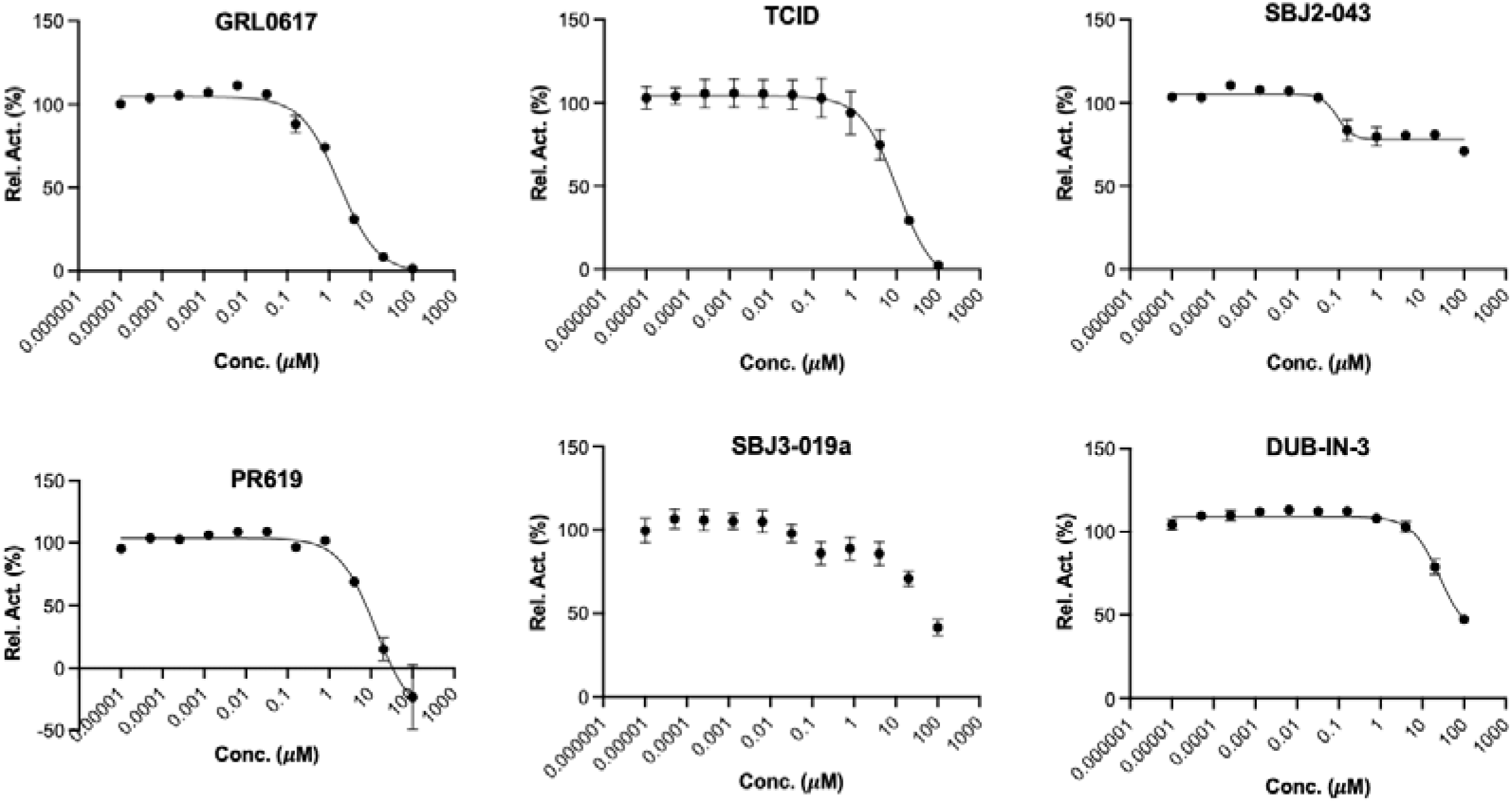
IC_50_ assays for 6 deubiquitinase inhibitors on their inhibition of 20 nM PLpro using 5 μM Ub-AMC as substrate. Experiments at different conditions were performed in triplicates.

To understand how some inhibitors may interact with PLpro, we conducted molecular docking using Autodock Vina. As a starting point for docking, we used the crystal structure of PLpro bound with GRL0617 (PDB ID: 7CMD).^27, 29^ GRL0617 and water molecules were removed from the active site to prepare PLpro for the docking analysis. To simplify our analysis, we limited docking around the active site. A binding pocket was defined based on the known residues of the S3/S4 binding pocket site of PLpro. 3D Conformers of selected compounds were generated using OpenBabel. We validated the docking protocol by conducting re-docking of GRL0617. Our docking results suggested a very similar binding mode of GRL0617 with PLpro as the co-crystal structure (Figure 7A). GRL0617 formed two hydrogen bonds between its amides and Asp164 and Tyr268, one hydrogen bond between its oxygen atom and Gln269. In addition, the naphthalene group of GRL0617 involves the hydrophobic interactions with aromatic residues Tyr264 and Tyr268. Other inhibitors ML364, LDN-57444, TCID and DUB-IN-3 were also docked against PLpro. The protein-ligand interactions and their detailed binding modes were illustrated in Figure 7. ML364 formed a hydrogen bond with the Asp164. Three additional hydrogen bonds with Arg166 were formed with ML364 (Figure 7B). LDN-57444, TCID, and DUB-IN-3 formed hydrogen bonds with Tyr268, Tyr264, and Arg166, respectively (Figure 7C-E). There is no strong correlation between calculated binding energies and determined IC_50_ values. This discrepancy may be explained by limited factors involved in the calculation that would lead to missing some of the potentially important interactions. Other contributing factors include potential covalent interactions with the enzyme. The crystallographic studies would be critical to understand the mechanism of inhibition of PLpro at atomic level and provide valuable insights in developing PLpro inhibitors with improved properties.

**Figure 7.**
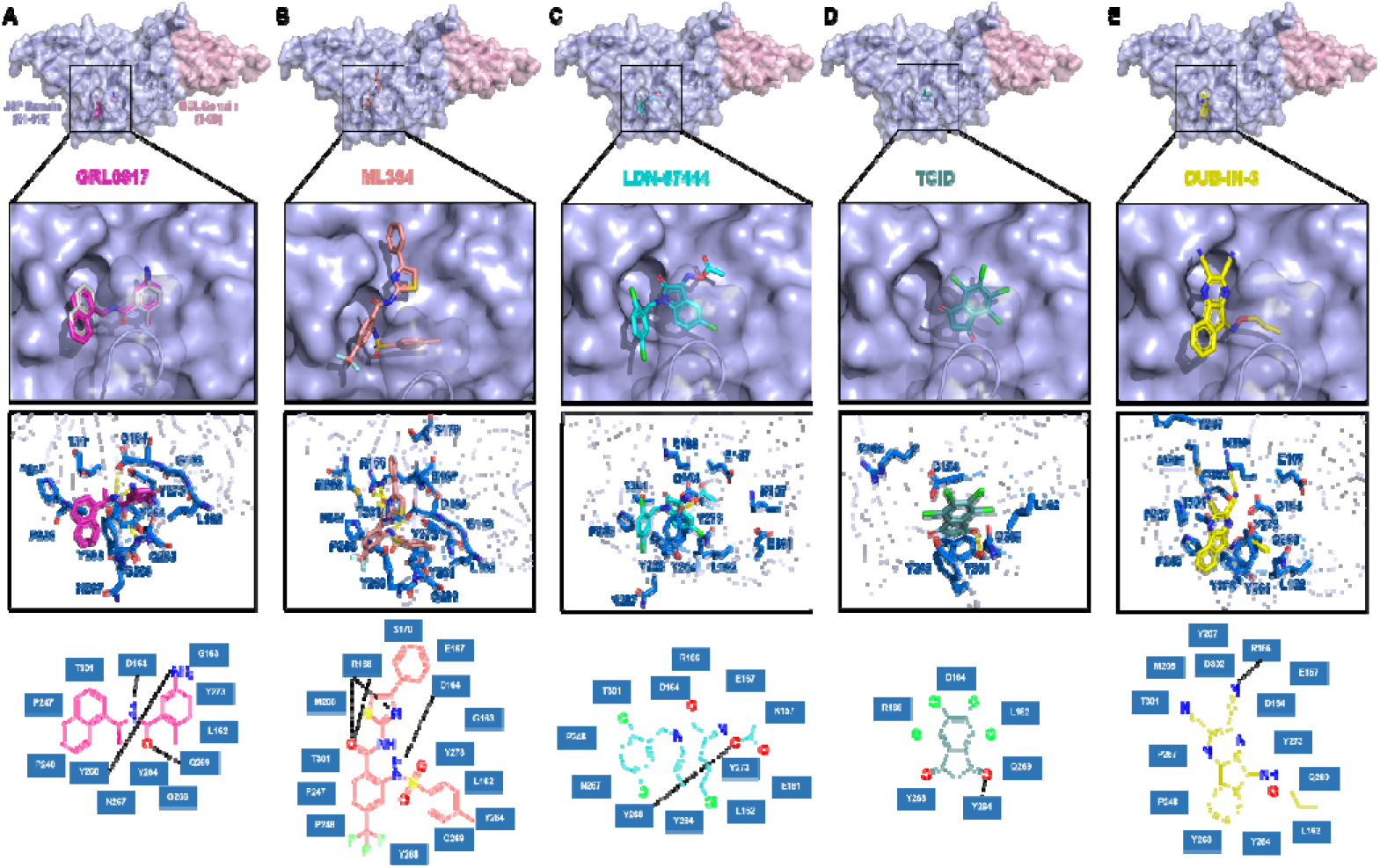
The top binding modes of the selected compounds along with their corresponding interactions within the active site of PLpro.

## CONCLUSION

Since 2003, there have been three coronavirus disease outbreaks. Many researchers have predicted that additional coronavirus diseases will emerge with higher frequency. For both combating the current pandemic and preparing to contain future coronavirus disease outbreaks, it is imperative to develop small molecule antivirals that may be applied generally to inhibit coronaviruses. Due to its conserveness among coronaviruses, PLpro is an attractive drug target for developing broad-spectrum antivirals. In this study, we experimentally characterized 33 deubiquitinase inhibitors and 37 cysteine protease inhibitors on their inhibition of PLpro from SARS-CoV-2. From this study, we identified seven molecules that potently inhibit PLpro with an IC_50_ value below 60 μM when Z-LRGG-AMC was used a substrate. Five inhibitors GRL0617, SJB2-043, TCID, SJB3-019A, and PR-619 have an IC_50_ value below 10 μM. Interestingly, SBJ2-043 can only partially inhibit PLpro but has an outstanding IC_50_ value of 0.56 μM. When Ub-AMC was used as a substrate, an even lower IC_50_ value of 0.091 μM was determined. SJB2-043 might likely bind to an allosteric site of PLpro to exert its inhibition effect. As a pilot study, the current work indicates that drug repurposing for COVID-19 by targeting PLpro holds promises but in-depth investigation of these inhibitors in their mechanisms of action will likely reveal unique inhibition mechanisms of PLpro for more advanced development of potent PLpro inhibitors.

## MATERIALS AND METHODS

### Chemicals

The deubiquitinase inhibitor library and cysteine protease library were purchased from MedChemExpress. Two other deubiquitinase inhibitors, C527 and VLX1579, were purchased from Cayman Chemicals. The PLpro substrate, Z-LRGG-AMC, was purchased from Cayman Chemicals.

### In-silico Molecular Docking Study

#### Protein Preparation

The crystal structure of the SARS-CoV-2 PLpro in complex with inhibitor GRL0617 (PDB ID : 7CMD) was obtained from the RCSB Protein Data Bank (https://www.rcsb.org/). Only chain A of the homo-tetrameric structure was used in our docking study. The cognate ligand was extracted from the structure, which would be applied in the following redocking test. In AutoDockTools-1.5.7, water molecules were deleted and polar hydrogens were added to the structure. Finally, the prepared protein structure was converted into a PDBQT file for further docking studies.

#### Ligand Preparation

Compounds in SDF format were converted into PDB files using OpenBabel-3.1.1. These PDB files were further converted into PDBQT files via AutoDockTools-1.5.7 with default options.

#### Docking Parameters and Method

In addition to receptor and ligand preparations, AutoDockTools-1.5.7 was also used for grid parameter setting. The cognate ligand of crystal structure 7CMD suggested the inhibitor binding site. A grid box with dimensions 30 × 30 × 30 centered at the coordinates X = − 30.0, Y = − 12.0, and Z = − 30.0 was used to represent the search space. Then we applied AutoDock Vina docking protocol with options of 20 CPUs to use and maximum 100 binding modes to generate. Only the top binding affinities and binding modes were shown in this paper.

#### Docking Validation

We redocked the cognate ligand GRL0617 into its cognate receptor in order to verify the correctness of our docking method and parameters. The top row of Figure 4A showed the docked conformation in magenta and the crystal conformation in grey. The superposition of two conformations demonstrated the feasibility of our molecular docking method.

### Expression and Purification of PLpro

PLpro gene fragment was purchased from Integrated DNA Technologies (IDT), amplified with the primer pair that has XbaI and XhoI cutting sites. The amplified gene was inserted into pBAD-sfGFP vector. In the final construct (pBAD-sfGFP-PLpro), a hexahistidine tag is located at the N-terminus of sfGFP. A TEV cutting site is placed after the C-terminus of sfGFP which is followed by PLpro. Chemically competent Top10 E. coli cells were transformed with the pBAD-sfGFP-PLpro plasmid. Then, a single colony was picked, inoculated to 10 mL of 2YT with ampicillin and grew overnight at 37 °C. The overnight culture was inoculated to 1L of 2YT with 100 folds dilution and grew until OD 600 (optical density at 600 nm) reached 0.6. Then, the medium was cooled down to 20 °C and induced with 0.2 % L-arabinose for 24 hr at 20 °C. The cells were collected by centrifuging at 6000 rpm for 30 minsand resuspended in lysis buffer (20 mM Tris, 250 mM NaCl, 5% glycerol, 0.2% TritonX-100 and 1 mg/mL lysozyme at pH 7.8). The cells were lysed by sonication at 60% amplitude with 1 sec on 4 sec off for a total 5 min. The cell debris was removed by centrifuging at 16000 rpm for 30 min and the supernatant was loaded onto 3 mL of Ni-NTA resin (Genscript). A gravity flow Ni-NTA chromatography was performed. The Ni-NTA resin was washed with 10 times resin bed volume of wash buffer (20 mM Tris, 250 mM NaCl and 10 mM Imidazole at pH 7.8) and eluted with 20 mL elution buffer (20 mM Tris, 30 mM NaCl and 300 mM imidazole at pH 7.8). The buffer was changed to 20 mM Tris and 30 mM NaCl with 20% glycerol by using HiPrep 26/10 desalting column. Finally, the protein was aliquoted, flash frozen with liquid nitrogen and stored in − 80 °C freezer.

### Expression and Purification of Ub-G76C-6H

The Ub-G76C-6H was expressed according to the protocol described before.^24^ Briefly, an overnight culture was inoculated to 2YT medium with 100 times dilution. It was grown in 37 °C to reach OD 600 of 0.7−0.9 and the protein was expressed overnight at 18 °C by 1 mM IPTG induction. The cells were harvested by centrifugation and resuspended in lysis buffer (50 mM NaH_2_PO_4_, 500 mM NaCl, 5 mM imidazole, 1 mM TCEP, pH 7.8) for sonication, and then the lysate was removed by centrifugation (16000 rpm, 30 min at 4 °C). The supernatant was collected and acidified with 6 M HCl dropwise until it reached pH 2 then stirred for 5-10 min to precipitate impurities. The impurities were removed by centrifugation (10000 rpm, 30 min at 4 °C), and then the supernatant was neutralized to pH 7.8 using 6M NaOH. The supernatant was loaded onto Ni-NTA resin and washed with washing buffer (50 mM NaH_2_PO_4_, 500 mM NaCl, 25 mM imidazole, 1 mM TCEP at pH 7.8) and then eluted with elution buffer (washing buffer substituting 25 mM imidazole with 300 mM imidazo le). Finally, the elution was desalted into 50 mM ABC buffer using HiPrep Desalting column (GE Healthcare: Chicago, IL, USA) and lyophilized into 100 nmol per aliquot.

### Synthesis of Ub-AMC

Ub-AMC was synthesized according to the Activated Cysteine-directed Protein Ligation (ACPL) technique described before.^24^ 100 nmol of lyophilized Ub-G76C-6H was dissolved in 194.5 μL of 50 mM borate (pH 9) solution supplied with 0.5 μL 500 mM TCEP. It was then mixed with 300 μL of 40 mM Gly-AMC (dissolved in DMSO) to make a reaction solution with 24 mM Gly-AMC, 20 mM borate and 0.5 mM TCEP in a 60% DMSO solution. The reaction was initiated by adding 5 μL 500 mM NTCB (dissolved in DMSO) and the pH value was adjusted to 9.5 by 6 M NaOH. The reaction was performed at 37 °C for 16 h and terminated by desalting it to 50 mM ABC buffer with HiTrap Desalting column (GE Healthcare: Chicago, IL, USA). It was further purified using Ni-NTA resin and collected the flow-through to remove the side product, dehydroalanine, that we discovered during the ACPL reactions.^35^ The purified Ub-AMC was analyzed by Orbitrap ESI-MS, then flash frozen and stored in − 80 °C.

### Screening Assay

To have a preliminary test of inhibition ability of each drug, a screening assay was performed. 50 μL reaction solutions with 40 nM of PLpro and 400 μM of each drug (deubiquitinase inhibitors and cysteine protease inhibitors) were preincubated in PLpro assay buffer (20 mM Tris, 300 mM NaCl at pH 7.5) with 4 mM DTT in 37 °C for 30 min. It was then mixed with 50 μL solutions with 100 μM of Z-LRGG-AMC. The final assay solution had 200 μM of each drug, 20 nM PLpro, 2 mM DTT and 50 μM Z-LRGG-AMC. The fluorescence of AMC that was generated as a result of PLpro enzymatic activity was recorded with plate reader by using 380 nm as excitation wavelength and 440 nm as the emission wavelength. The initial slope of the fluorescence vs time graph for each drug was analyzed by calculating the slope of the curve between 0−10 min. The calculation was done by using linear regression analysis in GraphPad 8.0. Then, the initial slopes were normalized against reaction substitute drugs with DMSO.

### Inhibition Analysis

The inhibition assay was performed the same way as the screening assay except varying the drug concentration with 200 μM and 100 μM serially diluted 5 for 10 times to obtain the inhibition curve. All the reactions are performed as triplicates. The further calculation was done through GraphPad 8.0. Initial slopes were calculated from the slope of first 10 min fluorescence increasement as described above. It was then normalized against the control reaction that has DMSO only. The normalized PLPro activity against drug concentration was analyzed with the inhibition curve (three parameters) analysis option in GraphPad 8.0 to determine the IC_50_ values.

**Table 1.**
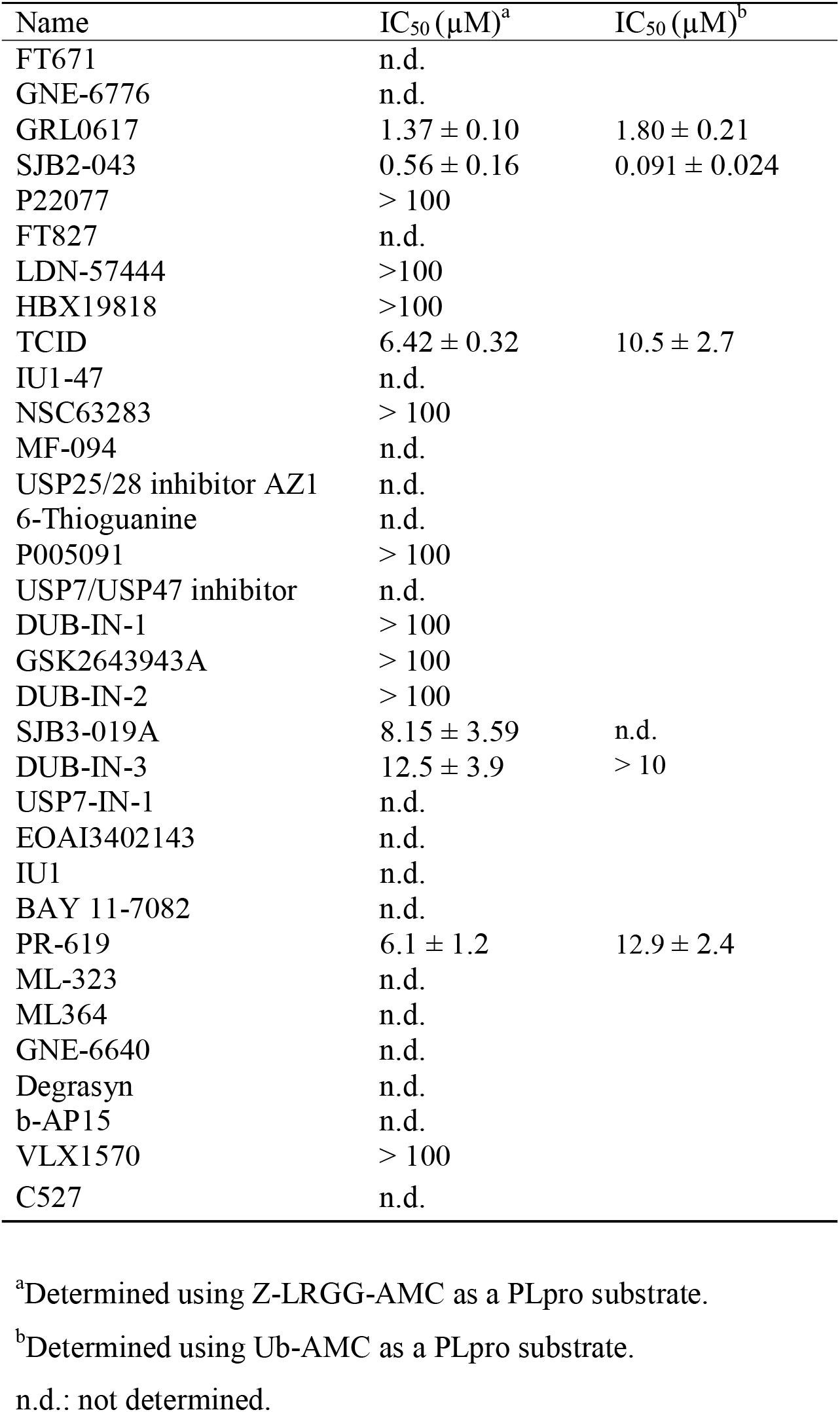
IC_50_ values of identified deubiquitinase inhibitors against PLpro.

**Table 2.** IC_50_ values of identified cysteine protease inhibitors against SC2PLpro.

| Name | IC<sub>50</sub> (μM) |
| --- | --- |
| NSC18058 | n.d. |
| Leupeptin | n.d. |
| HY-17541A | n.d. |
| HY-17541 | n.d. |
| PMSF | n.d. |
| EP1013 | n.d. |
| S130 | 35.0 ± 1.4 |
| Petesicatib | n.d. |
| Tauroursodeoxycholate | n.d. |
| Emricasan | n.d. |
| N-Ethylmaleimide | n.d. |
| CA-074 | n.d. |
| CA-074 methyl ester | n.d. |
| Z-VAD(OMe)-FMK | n.d. |
| Aloxistatin | n.d. |
| Q-VD-OPh | n.d. |
| H-Arg-4MβNA | n.d. |
| LY 3000328 | n.d. |
| BOC-D-FMK | n.d. |
| Z-DEVD-FMK | n.d. |
| E-64 | n.d. |
| Balicatib | n.d. |
| Odanacatib | n.d. |
| Belnacasan | n.d. |
| Calpetin | n.d. |
| E 64c | n.d. |
| 3-Epiursolic Acid | n.d. |
| MDL-28170 | n.d. |
| Chelidonic acid | n.d. |
| MG-101 | n.d. |
| Z-VAD-FMK | n.d. |
| OT-82 | n.d. |
| IKarisoside A | n.d. |
| Wedelolactone | n.d. |
| Phenoxodiol | n.d. |
| 2-Cyanopyrimidine | n.d. |
| Z-WEHD-FMK | n.d. |

## ACKNOWLEDGEMENTS

This work was supported in part by Welch Foundation (grant A-1715), Texas A&M Presidential Impact Fellowship Fund and Texas A&M X-Grant.

## REFERENCES

1. Gates, B., Responding to Covid-19 - A Once-in-a-Century Pandemic? N Engl J Med 2020, 382 (18), 1677–1679.

2. Morens, D. M.; Daszak, P.; Taubenberger, J. K., Escaping Pandora’s Box - Another Novel Coronavirus. N Engl J Med 2020, 382 (14), 1293–1295.

3. Morse, J. S.; Lalonde, T.; Xu, S.; Liu, W. R., Learning from the Past: Possible Urgent Prevention and Treatment Options for Severe Acute Respiratory Infections Caused by 2019-nCoV. Chembiochem 2020, 21 (5), 730–738.

4. Vatansever, E. C.; Yang, K. S.; Drelich, A. K.; Kratch, K. C.; Cho, C. C.; Kempaiah, K. R.; Hsu, J. C.; Mellott, D. M.; Xu, S.; Tseng, C. K.; Liu, W. R., Bepridil is potent against SARS-CoV-2 in vitro. Proc Natl Acad Sci U S A 2021, 118 (10).

5. Wang, Y.; Zhang, D.; Du, G.; Du, R.; Zhao, J.; Jin, Y.; Fu, S.; Gao, L.; Cheng, Z.; Lu, Q.; Hu, Y.; Luo, G.; Wang, K.; Lu, Y.; Li, H.; Wang, S.; Ruan, S.; Yang, C.; Mei, C.; Wang, Y.; Ding, D.; Wu, F.; Tang, X.; Ye, X.; Ye, Y.; Liu, B.; Yang, J.; Yin, W.; Wang, A.; Fan, G.; Zhou, F.; Liu, Z.; Gu, X.; Xu, J.; Shang, L.; Zhang, Y.; Cao, L.; Guo, T.; Wan, Y.; Qin, H.; Jiang, Y.; Jaki, T.; Hayden, F. G.; Horby, P. W.; Cao, B.; Wang, C. Remdesivir in adults with severe COVID-19: a randomised, double-blind, placebo-controlled, multicentre trial. 1474–547X (Electronic) 0140-6736 (Linking), May 16, 2020.

6. Lam, T. T.; Jia, N.; Zhang, Y. W.; Shum, M. H.; Jiang, J. F.; Zhu, H. C.; Tong, Y. G.; Shi, Y. X.; Ni, X. B.; Liao, Y. S.; Li, W. J.; Jiang, B. G.; Wei, W.; Yuan, T. T.; Zheng, K.; Cui, X. M.; Li, J.; Pei, G. Q.; Qiang, X.; Cheung, W. Y.; Li, L. F.; Sun, F. F.; Qin, S.; Huang, J. C.; Leung, G. M.; Holmes, E. C.; Hu, Y. L.; Guan, Y.; Cao, W. C., Identifying SARS-CoV-2-related coronaviruses in Malayan pangolins. Nature 2020, 583 (7815), 282–285.

7. Coronaviridae Study Group of the International Committee on Taxonomy of, V., The species Severe acute respiratory syndrome-related coronavirus: classifying 2019-nCoV and naming it SARS-CoV-2. Nat Microbiol 2020, 5 (4), 536–544.

8. Zhou, P.; Yang, X. L.; Wang, X. G.; Hu, B.; Zhang, L.; Zhang, W.; Si, H. R.; Zhu, Y.; Li, B.; Huang, C. L.; Chen, H. D.; Chen, J.; Luo, Y.; Guo, H.; Jiang, R. D.; Liu, M. Q.; Chen, Y.; Shen, X. R.; Wang, X.; Zheng, X. S.; Zhao, K.; Chen, Q. J.; Deng, F.; Liu, L. L.; Yan, B.; Zhan, F. X.; Wang, Y. Y.; Xiao, G. F.; Shi, Z. L., A pneumonia outbreak associated with a new coronavirus of probable bat origin. Nature 2020, 579 (7798), 270–273.

9. Phan, T., Genetic diversity and evolution of SARS-CoV-2. Infect Genet Evol 2020, 81, 104260.

10. Chen, Y. W.; Yiu, C. B.; Wong, K. Y., Prediction of the SARS-CoV-2 (2019-nCoV) 3C-like protease (3CL (pro)) structure: virtual screening reveals velpatasvir, ledipasvir, and other drug repurposing candidates. F1000Res 2020, 9, 129.

11. Naqvi, A. A. T.; Fatima, K.; Mohammad, T.; Fatima, U.; Singh, I. K.; Singh, A.; Atif, S. M.; Hariprasad, G.; Hasan, G. M.; Hassan, M. I., Insights into SARS-CoV-2 genome, structure, evolution, pathogenesis and therapies: Structural genomics approach. Biochim Biophys Acta Mol Basis Dis 2020, 1866 (10), 165878.

12. Shin, D.; Mukherjee, R.; Grewe, D.; Bojkova, D.; Baek, K.; Bhattacharya, A.; Schulz, L.; Widera, M.; Mehdipour, A. R.; Tascher, G.; Geurink, P. P.; Wilhelm, A.; van der Heden van Noort, G. J.; Ovaa, H.; Muller, S.; Knobeloch, K. P.; Rajalingam, K.; Schulman, B. A.; Cinatl, J.; Hummer, G.; Ciesek, S.; Dikic, I., Papain-like protease regulates SARS-CoV-2 viral spread and innate immunity. Nature 2020, 587 (7835), 657–662.

13. Klemm, T.; Ebert, G.; Calleja, D. J.; Allison, C. C.; Richardson, L. W.; Bernardini, J. P.; Lu, B. G.; Kuchel, N. W.; Grohmann, C.; Shibata, Y.; Gan, Z. Y.; Cooney, J. P.; Doerflinger, M.; Au, A. E.; Blackmore, T. R.; van der Heden van Noort, G. J.; Geurink, P. P.; Ovaa, H.; Newman, J.; Riboldi-Tunnicliffe, A.; Czabotar, P. E.; Mitchell, J. P.; Feltham, R.; Lechtenberg, B. C.; Lowes, K. N.; Dewson, G.; Pellegrini, M.; Lessene, G.; Komander, D., Mechanism and inhibition of the papain-like protease, PLpro, of SARS-CoV-2. EMBO J 2020, 39 (18), e106275.

14. Baez-Santos, Y. M.; St John, S. E.; Mesecar, A. D., The SARS-coronavirus papain-like protease: structure, function and inhibition by designed antiviral compounds. Antiviral Res 2015, 115, 21–38.

15. McClain, C. B.; Vabret, N., SARS-CoV-2: the many pros of targeting PLpro. Signal Transduct Target Ther 2020, 5 (1), 223.

16. Totura, A. L.; Bavari, S., Broad-spectrum coronavirus antiviral drug discovery. Expert Opin Drug Discov 2019, 14 (4), 397–412.

17. Liu, C.; Zhou, Q.; Li, Y.; Garner, L. V.; Watkins, S. P.; Carter, L. J.; Smoot, J.; Gregg, A. C.; Daniels, A. D.; Jervey, S.; Albaiu, D., Research and Development on Therapeutic Agents and Vaccines for COVID-19 and Related Human Coronavirus Diseases. ACS Cent Sci 2020, 6 (3), 315–331.

18. Petushkova, A. I.; Zamyatnin, A. A., Jr., Papain-Like Proteases as Coronaviral Drug Targets: Current Inhibitors, Opportunities, and Limitations. Pharmaceuticals (Basel) 2020, 13 (10).

19. Rut, W.; Lv, Z.; Zmudzinski, M.; Patchett, S.; Nayak, D.; Snipas, S. J.; El Oualid, F.; Huang, T. T.; Bekes, M.; Drag, M.; Olsen, S. K., Activity profiling and crystal structures of inhibitor-bound SARS-CoV-2 papain-like protease: A framework for anti-COVID-19 drug design. Sci Adv 2020, 6 (42), eabd4596.

20. Lei, J.; Kusov, Y.; Hilgenfeld, R., Nsp3 of coronaviruses: Structures and functions of a large multi-domain protein. Antiviral Res 2018, 149, 58–74.

21. Lindner, H. A.; Fotouhi-Ardakani, N.; Lytvyn, V.; Lachance, P.; Sulea, T.; Menard, R., The papain-like protease from the severe acute respiratory syndrome coronavirus is a deubiquitinating enzyme. J Virol 2005, 79 (24), 15199–208.

22. Ratia, K.; Saikatendu, K. S.; Santarsiero, B. D.; Barretto, N.; Baker, S. C.; Stevens, R. C.; Mesecar, A. D., Severe acute respiratory syndrome coronavirus papain-like protease: structure of a viral deubiquitinating enzyme. Proc Natl Acad Sci U S A 2006, 103 (15), 5717–22.

23. Liu, M.; Wang, B.; Wang, F.; Yang, Z.; Gao, D.; Zhang, C.; Ma, L.; Yu, X., Soluble expression of single-chain variable fragment (scFv) in Escherichia coli using superfolder green fluorescent protein as fusion partner. Appl Microbiol Biotechnol 2019, 103 (15), 6071–6079.

24. Qiao, Y.; Yu, G.; Kratch, K. C.; Wang, X. A.; Wang, W. W.; Leeuwon, S. Z.; Xu, S.; Morse, J. S.; Liu, W. R., Expressed Protein Ligation without Intein. J Am Chem Soc 2020, 142 (15), 7047–7054.

25. Ratia, K.; Pegan, S.; Takayama, J.; Sleeman, K.; Coughlin, M.; Baliji, S.; Chaudhuri, R.; Fu, W.; Prabhakar, B. S.; Johnson, M. E.; Baker, S. C.; Ghosh, A. K.; Mesecar, A. D., A noncovalent class of papain-like protease/deubiquitinase inhibitors blocks SARS virus replication. Proc Natl Acad Sci U S A 2008, 105 (42), 16119–24.

26. Baez-Santos, Y. M.; Barraza, S. J.; Wilson, M. W.; Agius, M. P.; Mielech, A. M.; Davis, N. M.; Baker, S. C.; Larsen, S. D.; Mesecar, A. D., X-ray structural and biological evaluation of a series of potent and highly selective inhibitors of human coronavirus papain-like proteases. J Med Chem 2014, 57 (6), 2393–412.

27. Fu, Z.; Huang, B.; Tang, J.; Liu, S.; Liu, M.; Ye, Y.; Liu, Z.; Xiong, Y.; Zhu, W.; Cao, D.; Li, J.; Niu, X.; Zhou, H.; Zhao, Y. J.; Zhang, G.; Huang, H., The complex structure of GRL0617 and SARS-CoV-2 PLpro reveals a hot spot for antiviral drug discovery. Nat Commun 2021, 12 (1), 488.

28. Ratia, K.; Pegan, S.; Takayama, J.; Sleeman, K.; Coughlin, M.; Baliji, S.; Chaudhuri, R.; Fu, W.; Prabhakar, B. S.; Johnson, M. E.; Baker, S. C.; Ghosh, A. K.; Mesecar, A. D., A noncovalent class of papain-like protease/deubiquitinase inhibitors blocks SARS virus replication. Proceedings of the National Academy of Sciences 2008, 105 (42), 16119–16124.

29. Gao, X.; Qin, B.; Chen, P.; Zhu, K.; Hou, P.; Wojdyla, J. A.; Wang, M.; Cui, S., Crystal structure of SARS-CoV-2 papain-like protease. Acta Pharm Sin B 2021, 11 (1), 237–245.

30. Liu, Y.; Lashuel, H. A.; Choi, S.; Xing, X.; Case, A.; Ni, J.; Yeh, L. A.; Cuny, G. D.; Stein, R. L.; Lansbury, P. T., Jr., Discovery of inhibitors that elucidate the role of UCH-L1 activity in the H1299 lung cancer cell line. Chem Biol 2003, 10 (9), 837–46.

31. Altun, M.; Kramer, H. B.; Willems, L. I.; McDermott, J. L.; Leach, C. A.; Goldenberg, S. J.; Kumar, K. G.; Konietzny, R.; Fischer, R.; Kogan, E.; Mackeen, M. M.; McGouran, J.; Khoronenkova, S. V.; Parsons, J. L.; Dianov, G. L.; Nicholson, B.; Kessler, B. M., Activity-based chemical proteomics accelerates inhibitor development for deubiquitylating enzymes. Chem Biol 2011, 18 (11), 1401–12.

32. Fu, Y.; Hong, L.; Xu, J.; Zhong, G.; Gu, Q.; Gu, Q.; Guan, Y.; Zheng, X.; Dai, Q.; Luo, X.; Liu, C.; Huang, Z.; Yin, X. M.; Liu, P.; Li, M., Discovery of a small molecule targeting autophagy via ATG4B inhibition and cell death of colorectal cancer cells in vitro and in vivo. Autophagy 2019, 15 (2), 295–311.

33. Mistry, H.; Hsieh, G.; Buhrlage, S. J.; Huang, M.; Park, E.; Cuny, G. D.; Galinsky, I.; Stone, R. M.; Gray, N. S.; D’Andrea, A. D.; Parmar, K., Small-molecule inhibitors of USP1 target ID1 degradation in leukemic cells. Mol Cancer Ther 2013, 12 (12), 2651–62.

34. Colombo, M.; Vallese, S.; Peretto, I.; Jacq, X.; Rain, J. C.; Colland, F.; Guedat, P., Synthesis and biological evaluation of 9-oxo-9H-indeno[1,2-b]pyrazine-2,3-dicarbonitrile analogues as potential inhibitors of deubiquitinating enzymes. Chemmedchem 2010, 5 (4), 552–8.

35. Qiao, Y.; Yu, G.; Leeuwon, S. Z.; Liu, W. R., Site-specific conversion of cysteine in a protein to dehydroalanine using 2-nitro-5-thiocyanatobenzoic acid. Molecules 2021, 26 (9), 2619.

